# CAD-C reveals centromere pairing and near-perfect alignment of sister chromatids

**DOI:** 10.64898/2025.12.18.694906

**Authors:** Axel Delamarre, Maurane Reveil, David Pintor Pichon, Michael A. Boemo, Iestyn Whitehouse

## Abstract

Three-dimensional (3D) genome organization plays a central role in gene regulation, chromatin folding, and genome stability. Although chromosome-conformation capture (3C)–derived methods have revolutionized our understanding of genome architecture, most remain limited in resolution, in their capacity to detect multiway interactions and in their ability to distinguish sister chromatids. Here, we present CAD-C, a new chromatin-conformation capture strategy that uses Caspase-Activated DNase (CAD) to fragment chromatin. Fragmentation of chromatin to the nucleosome level by CAD digestion substantially enhances proximity ligation, enabling formation of multi-nucleosome ligation products. Nanopore sequencing of these long DNA molecules allows reconstruction of chromatin fiber connectivity and 3D contact maps with single nucleosome resolution. Importantly, CAD-C is able to identify sister-chromatid interactions at high resolution which reveals that centromeres are closely paired and that cohesin maintains sister chromatids in near perfect alignment where the same nucleosomes are associated across sisters. Such precise alignment has significant implications for chromosome structure and the mechanisms by which cohesion is established.

## INTRODUCTION

In eukaryotes, genome duplication produces two sister chromatids that must remain physically attached until mitosis to ensure accurate chromosome segregation^1,2^. The connection of sister chromatids, known as cohesion, is established in S-phase by the cohesin complex; a conserved SMC (Structural Maintenance of Chromosomes) ring-shaped complex capable of physically entrapping two molecules of DNA^3^. Beyond its canonical role in cohesion, cohesin also contributes to the spatial organization of chromatin during interphase by forming loops that shape chromatin domains^4–7^.

Over the past two decades, our understanding of 3D genome architecture has been revolutionized by chromosome conformation capture (3C) methods such as Hi-C ^6,8,9^, which notably revealed the existence of chromatin compartments, topologically associating domains (TADs) and cohesin mediated chromatin loops. However, sequencing-based approaches face an inherent ambiguity when assaying mitotic chromosomes: because sister chromatids share identical sequences, interactions cannot easily be assigned to *cis* (within the same chromatid) or *trans* (between sisters) contacts. This issue is even more challenging in diploid genomes, where homologous chromosomes also share sequence identity.

To overcome this limitation, specialized variations of Hi-C have been developed to distinguish sister chromatid interactions. In yeast, Sister-C relies on the incorporation of the thymidine analog BrdU during replication and then uses a treatment with Hoechst and UV to selectively degrade the newly synthesized strand; ligation of the remaining parental strand is subsequently used to infer *cis* versus *trans*-sister contacts ^10^. Sister-C revealed that sister chromatids are typically well aligned at centromeric regions, but are generally loosely attached at cohesin enriched regions, with sister chromatids potentially paired across adjacent cohesin enriched sites – meaning that sister chromatids are not perfectly aligned. Although effective, this strategy partially destroys the substrate, which can reduce data yield and potentially introduce biases. An alternative approach, scs-Hi-C (sister-chromatid-sensitive Hi-C), incorporates 4-thio-thymidine during replication and applies a nucleoside conversion step to transform 4sdT→dC specifically on nascent strands, enabling direct sister discrimination without strand degradation ^11,12^. Applied in metazoans, scs-Hi-C could detect alignment of sister chromatids at cohesin and CTCF sites while observing a looser connection in regions depleted of cohesins. Scs-Hi-C also detected variations in alignment along the genome, tighter in silenced chromatin and looser over expressed genomic regions. A recent iteration combined Sister-C with Pore-C into Sister-pore-C ^13^; which detect BrdU incorporation on Oxford Nanopore sequencing improving the yield of scs-HI-C. A shared limitation of these strategies, however, is their reliance on restriction enzyme–based Hi-C that, coupled to the sparsity of the thymidine analogue incorporation and detection, limits the resolution of *trans* interaction from a few to hundreds of kilobases.

Historically, the assessment of sister chromatid cohesion has been achieved through fluorescence microscopy using a LacO arrays bound by LacI fused to GFP ^14^. In G2 phase, this imaging approach detects a single fluorescent focus when sisters are cohesed, while cohesin depletion leads to the appearance of two separated foci ^1,2^. Although this assay demonstrates that sister chromatids are held together below the resolution limit of conventional microscopy, it provides limited information about the sister-chromatid interactions. Furthermore, this approach is restricted to monitoring only a few genomic sites per cell, precluding genome-wide analysis.

The introduction of Micro-C, which uses micrococcal nuclease (MNase) to digest chromatin down to the nucleosome level, has largely overcome the constraints of using restriction endonucleases in Hi-C^15,16^. Yet, Micro-C–based approaches are limited by DNA ligation inefficiency, typically joining two nucleosomes to yield di-nucleosome products rather than extended chains of ligated nucleosomes that would be expected if most ends of nucleosomal DNA are competent for ligation. Such inefficiency restricts the direct detection of multiway interactions and, more generally, leaves an incomplete view of 3D organization when more than two partners are involved, for example when multiple cohesin sites clusters to form hubs.

To overcome limitations of pairwise interaction mapping, several strategies have been developed. Within the 3C framework, Pore-C sequences Hi-C products on Nanopore, capturing multiway contacts but remaining limited by restriction-enzyme fragmentation and its resolution limits^17,18^. In parallel, proximity-ligation-independent methods such as GAM^19^, SPRITE^20^, ChIA-Drop^21^, and more recently, PCP^22^ detect multiway interactions directly. While powerful, these approaches are generally more complex and costly to implement than classical 3C assays, and among them only PCP currently achieves nucleosome-resolution 3D mapping.

How nucleosomes are arranged and interact with each other along the chromatin fiber remains incompletely understood. Historical models supported by EM imaging suggest a 30nm fiber organization with an attractive helical structure, where alternating nucleosomes (n and n+2) stack together^23–26^. Consistent with these structures, Micro-C can detect nucleosome interactions compatible with stacking in a helix^15,27^. More recently, a dynamic picture of chromatin fiber organization based on clutches of nucleosomes has been observed with high-resolution microscopy such as chromEMT and STORM^26,28^. With such imaging, nucleosomes were generally found to exist in less ordered conformations with small regions adopting disorganized zig-zag structures.

Here, we introduce an alternative method to map 3D genome organization. CAD-C uses Caspase-Activated DNase (CAD) to enzymatically digest crosslinked chromatin to the nucleosome level, followed by proximity ligation. Using *S. cerevisiae*, we show that CAD chromatin digestion drastically improves the efficiency of proximity ligation and enables the recovery of high-molecular-weight products resulting from the ligation of multiple nucleosomes together. The long DNA concatemers can be sequenced on the Oxford Nanopore platform which provides information to (i) detect pairwise and multiway 3D interactions at nucleosome resolution, map the connectivity of fragments originating from the same DNA fiber, thereby reconstructing single-molecule nucleosome organization beyond population averages, (iii) unambiguously detect interactions between sister chromatids.

We show that CAD-C alleviates key limitations of existing 3C approaches and expands the detection of chromatin architecture to reveal that sister chromatids are near perfectly aligned in G2 in *S. cerevisiae*.

## RESULTS

### Proximity ligation from CAD-digested chromatin

In Micro-C, crosslinked chromatin is fragmented to the nucleosome level using micrococcal nuclease (MNase), followed by proximity ligation. While nucleosomes typically ligate to their direct neighbors, they can also ligate to more distant nucleosomes along the linear genome. From these hybrid molecules, 3D genome interactions can be inferred and high-resolution contact maps computed^15,16^. While one might expect that this procedure results in the ligation of multiple nucleosomes in a chain, proximity ligation in Micro-C typically yields di-nucleosome-sized DNA products representing the ligation of only two mono-nucleosomes^15,29^. This inefficiency likely reflects the fact that MNase digestion is hard to control, and frequently leaves DNA ends that are difficult to modify and ligate^22^.

To increase proximity ligation efficiency, recent applications of Micro-C used detergent before proximity ligation to facilitate high efficiency end-repair and ligation of digested chromatin^30^. However, such treatment preceding proximity ligation raise concern that the integrity of chromatin organization may be disrupted.

In a recent study from this lab, we used CAD (Caspase-Activated DNase) as an alternative to MNase to fragment chromatin in an orthogonal approach to map 3D chromatin organization (PCP)^22^. We found that CAD does not typically over-digest nucleosomal DNA and leaves DNA-ends that are more amenable to end-repair and subsequent ligation in chromatin, reflecting that CAD is well suited to controlled chromatin digestion^31–33^. Based on this observation, we reasoned that CAD digestion would improve the efficiency of proximity ligation of nucleosomes which would allow 3D genome interactions to be mapped. To facilitate bioinformatic analysis and concatemer splitting, we chose to add short DNA linker sequences to the ends of nucleosomal DNA following CAD digestion; complementarity between linkers was then generated by base excision allowing proximity ligation of nearby nucleosomes (Fig.1a). Using CAD to enzymatically digest the crosslinked chromatin substantially increases ligation efficiency relative to MNase and yields high-molecular-weight products comprising multiple, concatenated, nucleosomes (Fig.1a–b), with an average size of ∼1.3 kb (Fig.1c).

**Figure 1:**
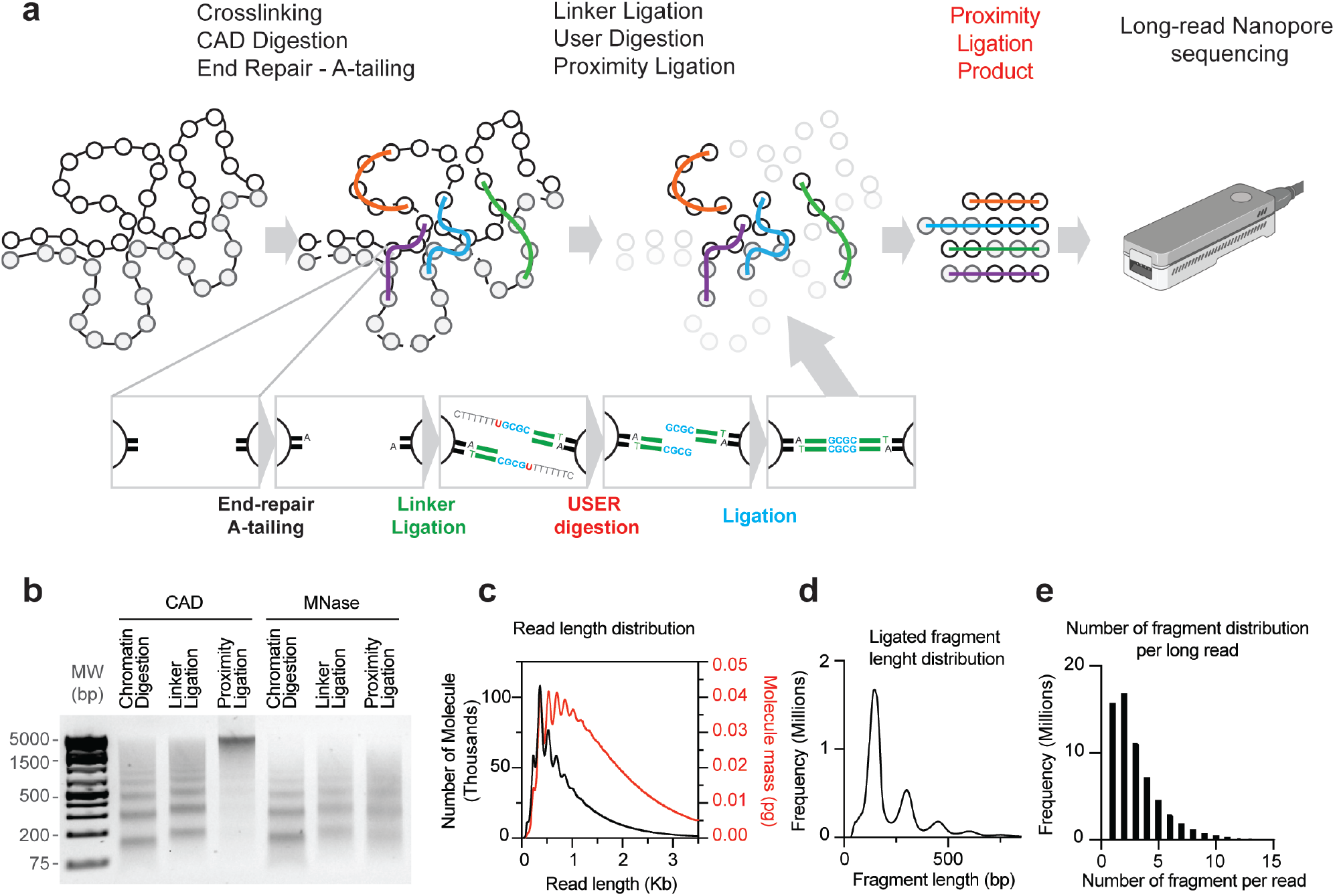
Principle of the CAD-C method: **a**) Schematic of the CAD-C. Crosslinked chromatin is digested using CAD to the nucleosome level. DNA ends are repaired, A-tailed and ligated to a DNA linker. DNA linkers are then activated by uracil base excision allowing proximity ligation to take place. CAD-C successfully ligates multiple nucleosomes in high molecular weight molecules that can then be sequenced by nanopore. **b**) Agarose gel of purified DNA species with control DNA sample after enzymatic digestion, linker ligation and proximity ligation. The efficiency of the reaction is higher when using CAD versus Micrococcal Nuclease (MNase) for chromatin enzymatic digestion. **c**) Read length distribution shown as number of molecules (black) and DNA mass (red) for comparison with b. **d**) Sequenced fragment length distribution after read splitting recapitulates the CAD chromatin digestion distribution. **e**) Distribution of the number of fragments ligated in long reads.

Long DNA molecules are purified and sequenced on Oxford Nanopore without amplification. Detection of the linker sequence allows the unambiguous segmentation of each long read into its constituent fragments. As expected, the size distribution of the sequenced DNA inserts closely match the initial CAD fragmentation pattern observed by gel electrophoresis (Fig.1b,d). Genome coverage of the sequenced DNA is homogeneous (Fig.S1a). The number of chromatin fragments per long read has a broad distribution, with a mean of ∼4 fragments per read (Fig.1e).

### Chromatin fragment composition of long-reads

Basic inspection of the nucleosome concatemers generated by CAD-C revealed a complex ligation pattern. As an example, we focused on the *MPH1* gene and mapped the connectivity of chromatin fragments. Each read is represented by a single line with shaded blocks denoting the footprints of mapped nucleosomes within the read. The orientation and the position within the long read are reported as well (Fig.2a,b). The plot is centered on the +1 nucleosome and reveals that this nucleosome preferentially interacts with adjacent nucleosomes within the gene body (density, shown in green), but overall, the interactions are complex, and nucleosomes are not necessarily ligated in the same order or orientation with which they are present in the genome (zoomed inset on right). Typically, in gene bodies, MNase-seq profiles show well positioned nucleosomes at the 5′ and 3′ ends but show a characteristic “fuzzy” pattern over gene bodies^34,35^. Population-averaged MNase-seq profiles can’t distinguish between genuinely disordered chromatin and regular arrays whose phasing varies between cells. Leveraging proximity ligation, CAD-C can capture chains of nucleosomes from the same fiber, allowing single-molecule reconstruction of their positions and spacing. We plotted all long reads mapping to the coding region of *MPH1* (Fig.2c) where each line corresponds to an individual read, and each black box corresponds to a footprint (e.g. nucleosome) contained with each long read. To facilitate visualization, reads are aligned by their rightmost position. The plot shows consistent positioning and spacing of nucleosomes at 5′ and 3’ ends of the gene, while revealing that apparently “fuzzy” regions are composed of regularly spaced but slightly offset positioned nucleosomes from fiber-to-fiber, hence from cell-to-cell. This result is consistent with our previous observations ^22^. On single fibers, nucleosomes tend to be well spaced in groups of 3–4 before consistent spacing fades (Fig.2c). Moreover, convergent genes lacking an intervening NFR form continuous chromatin units that are indistinguishable from tandem genes in terms of nucleosome fiber organization (Fig.S2a). Together, CAD-C enables single-fiber nucleosome positioning comparable to other single molecule approaches such as PCP or Fiber-seq^36^.

**Figure 2:**
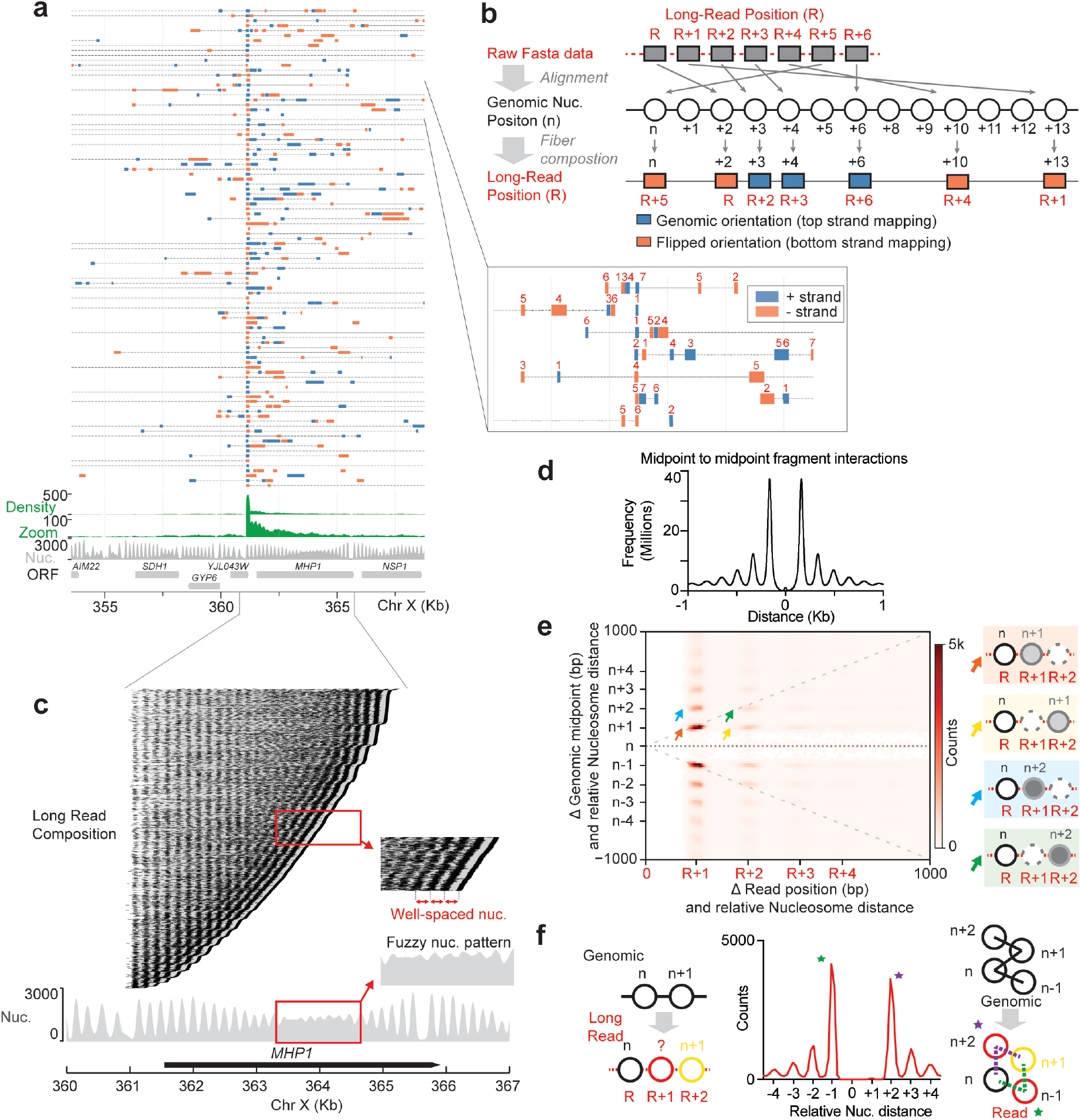
Long-read composition: **a**) A subset of long reads that contain a fragment aligning to the +1 nucleosome of the gene *MPH1* are shown. One long read is shown per line, fragments within each long read are represented as boxes denoting their size and orientation (+ strand in blue, − strand in red). The relative position that each mapped fragment is found within each long read is represented as a number in the zoomed-in panel. **b**) Schematic explaining data processing from raw data to long-read representation. **c**) Long read fragment composition over the gene *MPH1* in G1, each line corresponds to a long read and each black dot to a ligated chromatin fragment, e.g. nucleosomes. **d**) Midpoint to midpoint distance of pairwise ligation events of nucleosomes in each long read is shown. **e**) Heatmap of the relative position of a mapped fragment on a long read is plotted on the X-axis (R, R+1, etc…); the relative position on the genome is shown on the Y-axis (n, n+1, etc..). The orange arrow shows: R+1=n+1; the blue arrow shows: R+1=n+2; the green arrow shows: R+2=n+2 and the yellow arrow shows: R+2=n+1. The diagram on the right side represents the long read composition matching the color code of the arrows, colored ovals are nucleosomes. **f)** Schematic of R+2=n+1, a nucleosome intercalated between n and n+1 (i.e: yellow arrow in e.). Distribution of the relative genomic position for the intercalating nucleosome in shown in f: most of the intercalating nucleosomes are n-1 (green star) or n+2 (purple arrow). These configurations are compatible with a zig-zag organization or stacking.

### Pairwise ligation preferences of nucleosomes

To investigate whether there are any specific nucleosome-nucleosome interactions within the mapped data, we analyzed all pairwise interactions between mono-nucleosomes (21.7 million interactions) and measured ligation distances between midpoints of ligated nucleosomes. This analysis confirmed that nucleosomes ligate primarily to their direct neighbors (n+1 and n–1) (18.4%), followed by ∼2-fold less ligation to the next-nearest nucleosomes (n+2 and n–2) (8.5%) (Fig.2d). These preferences are robust and show minimal differences between highly expressed genes (1st sextile) and unexpressed genes (6th sextile) (Fig.S2b,c).

We then asked whether ligation to the following nucleosome reconstitutes native genomic orientation or favors flipped arrangements. This can be represented by distances between the endpoints of ligated fragments. Because we unambiguously map fragment sizes and orientations, we focused on ligation preferences of one extremity of a mono-nucleosome (Fig.S2d,e). Nucleosomes predominantly ligate to their immediate neighbors in the native genomic orientation: the distal end of a nucleosomal fragment most frequently ligates to the proximal end of the following nucleosome (Fig.S2d). The second most frequent class is split between ligation to the proximal end of the previous nucleosome (n–1) to the distal end of the following nucleosome (n+1). This ligation-preference pattern is robust across chromatin contexts, including the +1-nucleosome adjacent to nucleosome-free regions (NFRs), and is largely independent of transcription, being similar in expressed (1st sextile) and non-expressed genes (6th sextile) (Fig.S2f-i). Collectively, in CAD-C data, ligation between neighboring nucleosomes tend to reconstitute native order and orientation.

### Ligation preference of 3 consecutive mono-nucleosomes

To investigate how multiple nucleosomes arrange on the long read, we selected nanopore reads starting with at least three consecutive mono-nucleosomes (1.18 million, 1.4% of total read numbers) and plotted their relative genomic position (noted as n, n+1, n+2, etc…) against their position in read (noted as R, R+1, R+2 etc..) (Fig.2e,S2.j). If the proximity ligation product recapitulates the linear genomic organization, this plot should produce a diagonal where relative positions match (R+1 ∼ n+1, R+2 ∼ n+2 etc..). We observe a different result, where the first coordinate is strong at R+1 ∼ n+1, ∼8% of the selected data (Fig.2e, orange arrow, S2j), consistent with the pairwise analysis, but quickly fades, showing that proximity ligation tends not to simply follow the genomic linear organization across multiple nucleosomes. Instead, the third nucleosome on the long read (R+2) is as likely to be n+1 (∼4.4%, yellow arrow) as n+2 (∼4.6%, green arrow) (Fig.2e, S2j). This indicates that in some cases a distal nucleosome inserts between nucleosome neighbors. We selected these occurrences and plotted the relative genomic distance of the inserting nucleosomes (Fig.2f). This revealed that the intercalating nucleosomes mainly correspond to n-1 (Green star, i.e: n→**n-1**→ n+1; ∼0.7%) or n+2 (Purple star, i.e: n→ **n+2** → n+1 ∼0.7%) (Fig.2f, S2k). These ligation preferences are compatible with a local stacked or zig-zag organization of neighbor nucleosomes and while rare in the total population of ligation events, such zig-zag arrangements are as common as ligation in the linear genomic order G → G+1 → G+2 (∼1.4%). No differences in nucleosome repeat length (NRL) could be measured between pairwise direct neighbor nucleosomes, trimeric stacked nucleosomes or trimeric linear nucleosomes. (Fig.S2l). On longer fibers the contacts produce a diffuse signal indicating that no consistent, well organized nucleosome pattern is detected.

### Very high-resolution 3D genome interaction maps

CAD-C readily detects longer-range interactions than Micro-C, though not as extensive as proximity-ligation-independent approaches like PCP (Fig.S3a). CAD-C interaction maps are highly reproducible across replicates (Fig.S3b) and exhibit features consistent with established 3D genome architectures. At the whole-genome level, CAD-C detects strong inter-centromeric interactions corresponding to the Rabl configuration characteristic of yeast interphase nuclei (Fig.S3c). At intermediate scales, CAD-C consistently identifies chromatin interaction domains (CIDs) and their boundaries, correlating in location and strength with PCP data (Fig.S3d,e)^4,15,22^.

At very high resolution, CAD-C precisely maps nucleosome–nucleosome interactions: genes are separated by white stripes corresponding to NFRs (Fig.3a, S3f) and form their own small CIDs. Because fragment sizes are known, mapping midpoints further sharpens nucleosome-level interaction precision. Overall, CAD-C produces very high-resolution nucleosome interaction maps that outperform Micro-C and are comparable to PCP.

**Figure 3:**
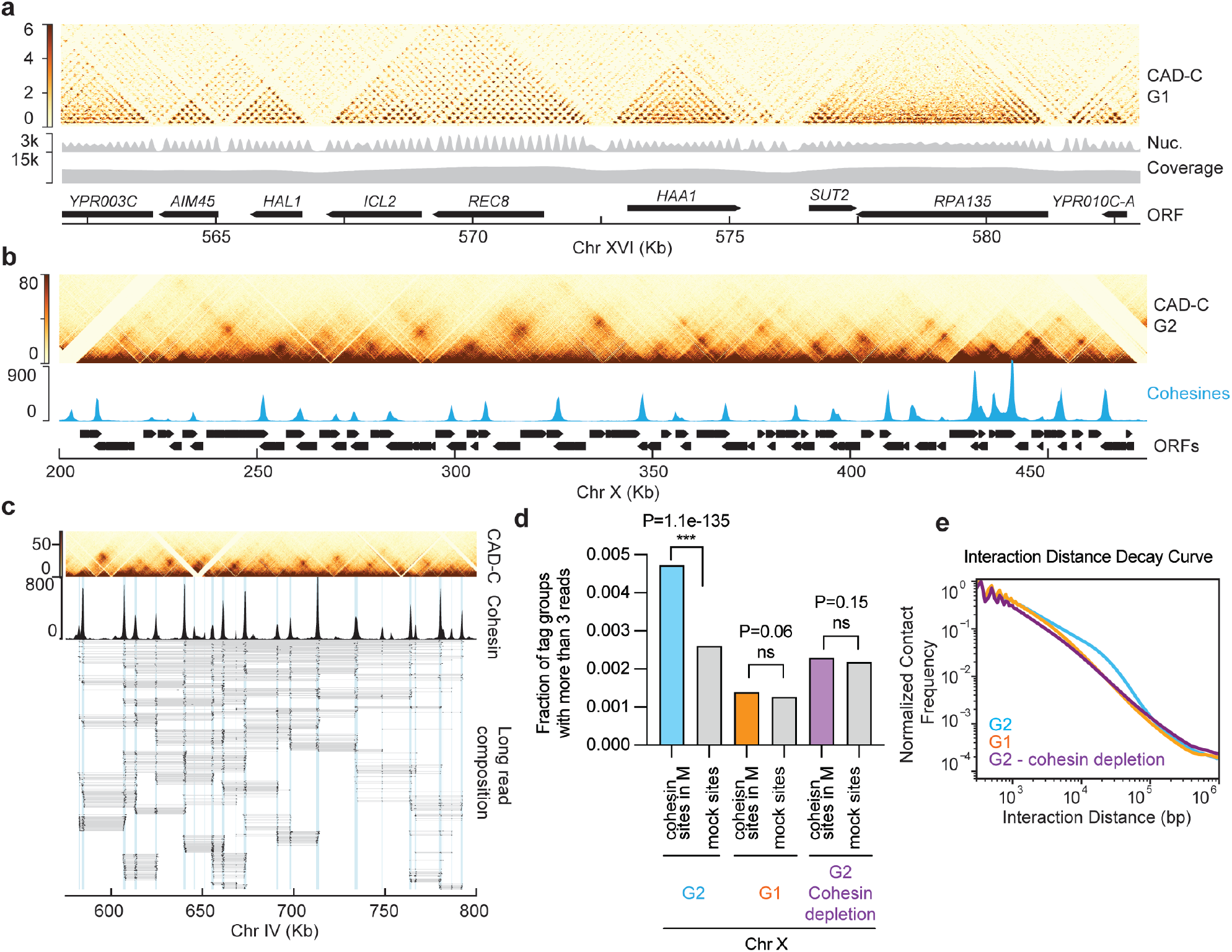
CAD-C produces high resolution 3D interaction maps and detects multiway interactions: **a**) Genome wide matrix of interaction produced by CAD-C in G1 at 10bp resolution. b) Genome wide map produced by CAD-C in G2 (Nocodazole arrest). Note the chromatin loop interactions between cohesin association sites. 250bp resolution. **c**) Long read composition connecting more than two cohesin accumulation sites. **d**) Quantification of the frequency multiway interactions with more than three fragments in different cohesin accumulation sites in G1, in G2 or in G2 after cohesin depletion. Note multi-way interacts are enhanced by cohesin and preferentially occur at cohesin loading sites. Fisher exact test is used to calculate p value. **e**) Interaction distance decay in CAD-C in G1, in G2 or in G2 after cohesin depletion.

### Multiway interaction detection

A hallmark of chromatin organization in Metaphase synchronized *S. cerevisiae* is the formation of cohesin-dependent loops along chromosome arms^4,5,22^. These loops are clearly visible in CAD-C (Fig.3b). As shown earlier, these loops are completely lost under depletion of the cohesin subunit Scc1^4,5^ (Fig.S4a).

Classical 3C methods only capture pairwise interactions, limiting direct detection of multiway contacts and their interpretation. For example, in cases where three genomic segments can be seen interacting in a pairwise manner, it is unclear whether apparent triplets reflect simultaneous tri-partite interactions or mutually exclusive pairwise events. Because CAD-C ligates chains of multiple nucleosomes, multiway interactions can be studied directly, overcoming this limitation. Using an orthogonal approach not relying on proximity ligation, we recently showed that cohesin forms hubs of chromatin loops^22^. CAD-C reproduces this result: multiple nucleosomes at distinct cohesin binding sites are found ligated together, confirming that cohesin loops tend to associate in space rather than being mutually exclusive (Fig.3c). We quantified the frequency of these long-range multiway interactions, observing enrichment at cohesin sites relative to segments between cohesin sites. As expected, these signals are reduced to background under cohesin depletion (Fig.3d). Accordingly, cohesin depletion disrupts both cohesin loops and multiway interactions (Fig.3d, S4a,b). CAD-C is therefore an alternative to PCP in the detection of multiway interactions at very high resolution.

### Detection of sister chromatid alignment at centromeres

Typical high-resolution interaction matrices display the relative frequency of pairwise interactions across genomic regions, with the interaction distance displayed on the y-axis perpendicular to the midpoint of their genomic coordinates. Thus, mapped interaction distances between the midpoints of adjacent nucleosomes are ∼165 bp (i.e. dyad to dyad distance, Fig4.a,b). Further exploring the interactions matrices produced by CAD-C, we noticed an unusual pattern at centromeres, where a strong signal was observed with an apparent interaction distance near 0 bp (Fig.4a,b green arrow). Remarkably this signal is visible across all centromeric regions in G2 but completely absent in G1 (Fig.4a-c).

One possibility that could explain this signal is if two small DNA fragments next to each other ligate at a very high frequency. To test this, we examined chromatin footprints at centromeres. Locus-specific footprint heatmaps derived from the CAD-C data show that centromeres yield fragments sizes slightly larger than canonical nucleosomes (Fig.4d,e), with clear asymmetry and a region of variability extending past the CDEIII element (Fig.4e). While this footprinting is in general agreement with MNase digestion profiles^37,38^, the pattern of CAD digestion provides a clearer picture of the nature of the centromeric nucleosome structure^39^.

Given that the majority of footprint sizes at the centromere are above 150bp (e.g Fig.4.d,e), the very short-range interactions in Figure 4b cannot be explained by the interactions of subnucleosomal fragments. We then considered the possibility that the short-range interactions were a result of contacts between sister chromatids, which are present in G2, but not in G1. Close inspection of individual nanopore reads covering centromeres reveals that the centromeric sequences can be present twice within a single long-read (Fig.4f, S5a). Given that CAD-C utilizes amplification-free, single molecule sequencing on Oxford Nanopore, the duplicated sequences cannot be explained by a PCR artefact. Further, the assay is conducted in a haploid strain and budding yeast utilizes “point centromeres” – defined by a single ∼120bp sequence per chromosome^40^, so duplicated centromeres should only be present following S-phase. Thus, the apparent ∼0bp interaction distance results from mapping the genomic distance between the midpoints of centromeric sequences present on two sister chromatids.

**Figure 4:**
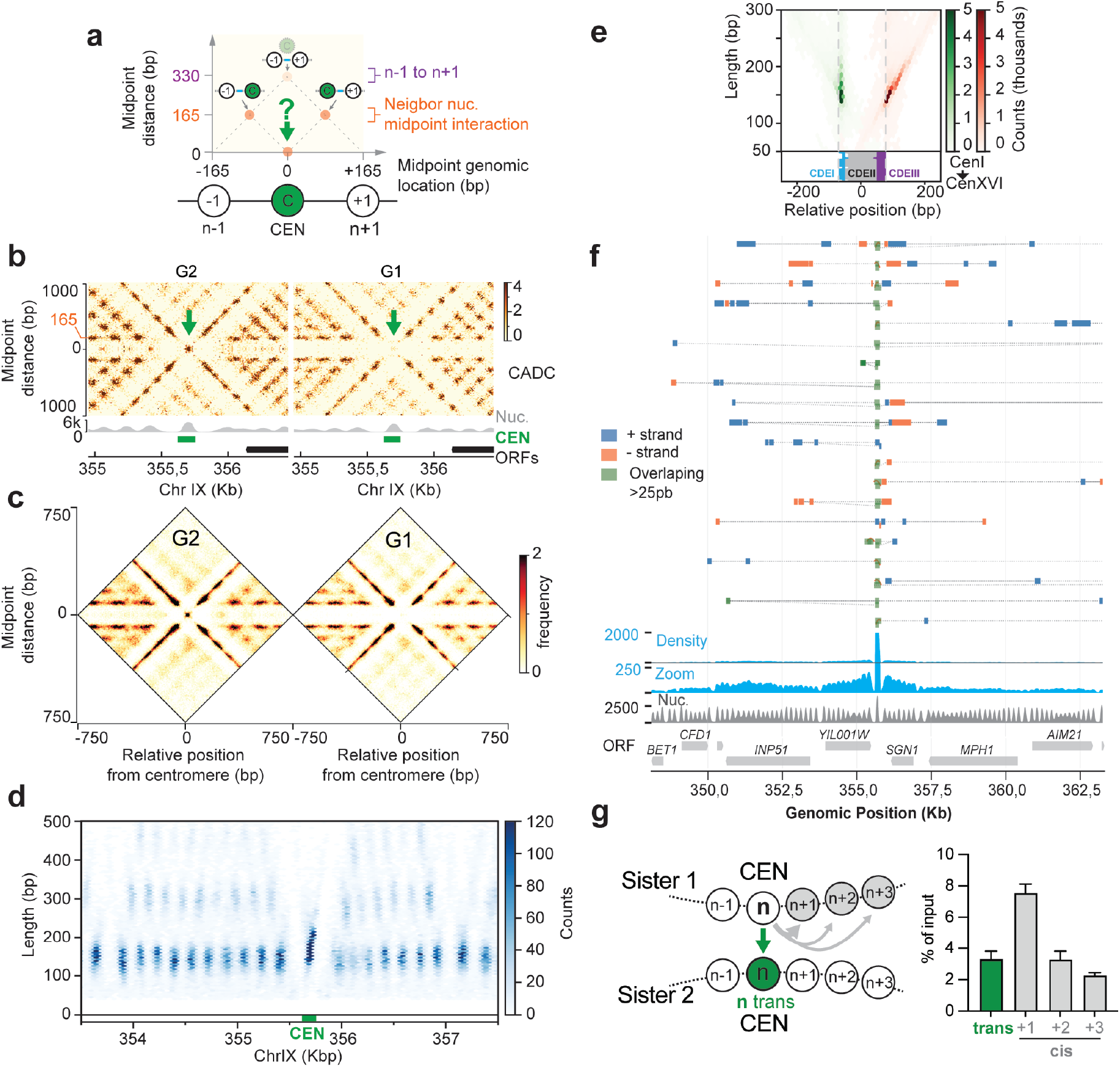
CAD-C detects sister chromatid alignment and proximity: **a**) Schematic describing the interpretation of interaction matrices at high resolution also highlighting the unusual pattern of interaction on the baseline of the plot with a midpoint distance of ∼0bp, green arrow. **b**) CAD-C interaction maps around the centromere of Chr IX in G1 (alpha-factor) and G2 (nocodazole) synchronized cells. Note the unusual interaction signal on the 0 axis in G2 absent in G1 (green arrow). The matrix was mirrored on the X axis for better visibility. **c**) as b but for all centromeres combined. **d**) Heatmap of mapped chromatin footprints around the centromere of Chr IX in G2, sequenced fragments are plotted according to their length (Y-axis) and position (X-axis), note the centromere has altered footprint. **e**) Metaplot of the left (green) and right (red) extremities of mapped DNA fragments for all 16 centromeres is shown relative to the centromere sequences. **f**) A fraction of the long reads from a G2 sample that contain two overlapping fragments aligning on the centromere of ChrIX are represented. One long read is shown per line, fragments are represented as boxes representing their size and orientation (+ strand in blue, − strand in red). Reads that overlap by at least 25pb are offset and highlighted in green. **g**) Quantification of the percentage of long reads containing the same centromere fragment twice vs the percentage of reads containing adjacent fragments corresponding to n+1, n+2 or n+3. Error bars: SD.

Next, we investigated the frequency at which the sister chromatids interact at the centromere. While absolute quantitation of interaction frequency is challenging, we can calculate the relative abundance of *cis* vs *trans* interactions of the centromeric nucleosome. For this we compared the frequency at which the centromeric nucleosome ligates to is direct neighbor nucleosomes (*in cis*) vs its sister centromere (*in trans*); remarkably, this revealed that centromeric nucleosomes interact with their sister centromere at a comparable frequency to which it interacts with nucleosomes directly adjacent *in cis* (Fig.4g). More specifically, the interaction frequency *in trans* matches that of the n+2 nucleosome which is separated from the centromere by less than <20 nm. Thus, sister centromeres are maintained in close proximity and may be effectively paired in G2.

### Sister-chromatid nucleosome ligation genome-wide

Beyond centromeres, interaction matrices revealed more extensive evidence of precise pairing between sister chromatids (Fig.5a, S6a). While this signal was less pronounced than at centromeres, it was clear that inter-sister interactions occurred at many sites along chromosomes. To quantitate interactions, we computed the frequency at which the same DNA sequence was represented twice within each nanopore read, such events should only occur when the same regions of the two sisters are ligated to each other. We then plotted the coverage of these overlapping regions to reveal the frequency of inter-sister contacts (Fig.5b, S6b). To facilitate visualization, this map was mirrored to represent the proximity between the sisters. This profile revealed peaks at centromeres and at cohesin accumulation sites defined by ChIP-seq^4,41^, with valleys between cohesin sites. Given that this assay can only detect perfect alignment between sisters, this profile indicates that sister chromatids are highly aligned with near perfect pairing at centromeres and at cohesin sites and slightly looser alignment between cohesin sites. Interactions between sisters at cohesin sites was reduced to ∼30% of that found at centromeres. Sister chromatin alignment does not decay as distance increase from the centromeres (Fig.S6c). As expected, the overlapping fragments profile is drastically reduced in the absence of cohesin in auxin-induced degradation of Scc1 or in G1 synchronized cells (Fig.5b,c, S6b).

**Figure 5:**
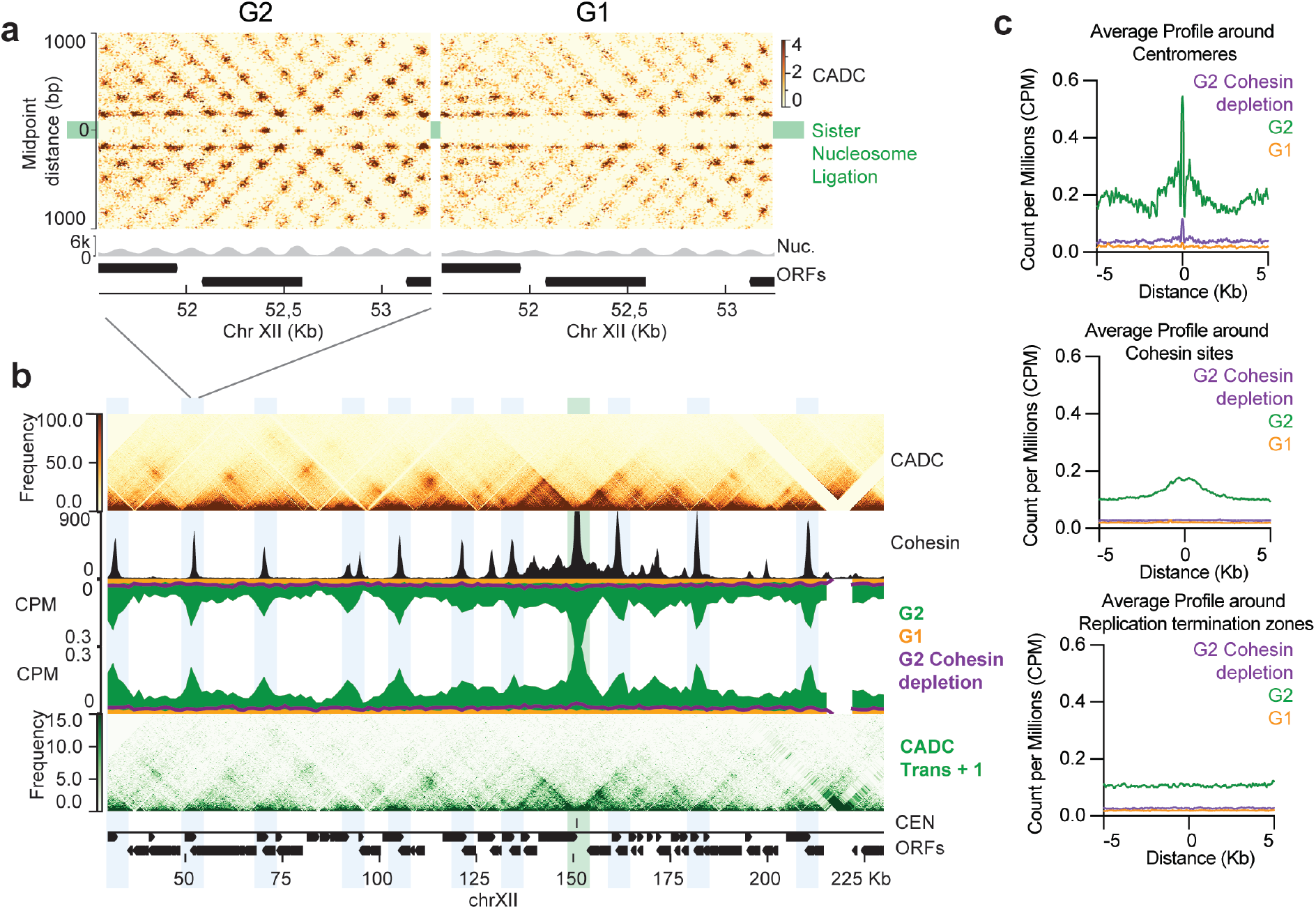
Genome-wide profile of sister nucleosome ligation: **a**) CAD-C interaction matrix around a cohesin accumulation site in G2, note the unusual signal on the 0 axis. This signal is absent from the G1 interaction matrix. The matrix was mirrored on the X axis for better visibility. **b**) Cad-C interaction matrix in G2 (top panel); cohesin ChIP dataset in G2 (black) (Costantino *et al*. 2020); mirrored sister chromatin ligation scores in G2 (green), G1 (orange) and with cohesin depletion in G2 (purple); interaction matrices based on long reads that simultaneously contain sister chromatid ligation fragments (lower panel). Green highlight represents centromere location. Blue highlights represent cohesin accumulation sites. **d**) Average profile of the sister-nucleosome ligation profile around centromeres, cohesin sites and replication termination sites in G2 (green), G1 (orange) and under cohesin depletion in G2 (purple).

The findings of close alignment apparently differ from results generated by Sister-C^10^, which proposed that sister chromatids are often not perfectly paired, but loosely aligned and frequently connected across adjacent cohesin sites (i.e: cohesin would pair cohesin site A on sister 1 to cohesin site B on sister 2). To reconcile our results, we considered a model in which cohesin tightly and near perfectly aligns sister chromatids while also forming multiway longer-range contacts with other loci (e.g. Fig.3c). Because Sister-C is insensitive to short range (less than 1.5kb) and multi-way contacts, an incomplete picture of paired sister chromatids is likely generated. To test this possibility, we selected long-reads that contain near perfectly aligned nucleosomes from both sisters, then plotted the interactions of the other nucleosomes within each of these reads (∼7.5% of all interactions); the resulting map clearly reproduces the global features of the full dataset (Fig.5b, lower panel in green), including long-range interactions and, prominently, cohesin accumulation sites. This indicates that nucleosomes are simultaneously aligned with their sister nucleosome while also part of long-range interactions.

### Sister-chromatid interaction detection by BrdU incorporation detection

While overlapping sequences within the same read provide a sensitive measure for sister chromatid interactions, the readout can only detect near perfect alignment between sisters. Thus, if the sisters are misaligned by just a few hundred base pairs then the present method would be incapable of resolving inter-sister contacts. To circumvent this problem, we leveraged nanopore’s ability to detect the incorporation of BrdU^42^ which can be used to distinguish sister chromatids based on the presence of the nucleotide analogue in a specific strand. Cells were arrested in G1, released into S-phase in the presence of BrdU, then arrested in G2 with nocodazole. The CAD-C protocol was conducted on these cells and the presence of BrdU was assessed in each mapped fragment. Theoretically, 50% of fragments should contain BrdU if the incorporation frequency is high and even across the genome. In practice we found that BrdU incorporation was variable and accurate detection was challenging on nucleosome-length reads. Comparing with a non-BrdU control dataset, we can nevertheless identify 4.4 million (∼1.5%) fragments as BrdU positive with stringent parameters (Fig.6a,S7a, cutoff 0.55). We then considered pairwise interactions where the two reads are BrdU positive; if the two ligated fragments have the same strandedness (+ to +, or − to −), the interaction is labeled *cis* (72% of the interactions) but if the strands are mixed (+ to −) the interaction is *trans* (28%) (Fig.6b,c). We then measured the distance between the two fragments in each interaction class (Fig.6c). As expected, *cis* interactions showed a strong dependence on genomic distance with adjacent nucleosomes along the same chromatin fiber preferentially interacting (ligating). Remarkably, we found a similar profile for interactions in *trans*, a result that agrees well with the analysis above but is nonetheless remarkable given that interactions between sister chromatids are not intrinsically constrained like *cis* interactions. Interactions in *trans* show a slightly broader profile than in *cis*, but the most prevalent *trans* interactions occur with a few hundred basepairs across the paired sisters. Across the genome, we found little variation in this pattern, including at differentially transcribed regions and cohesin enriched regions (Fig.7Sb-c).

**Figure 6:**
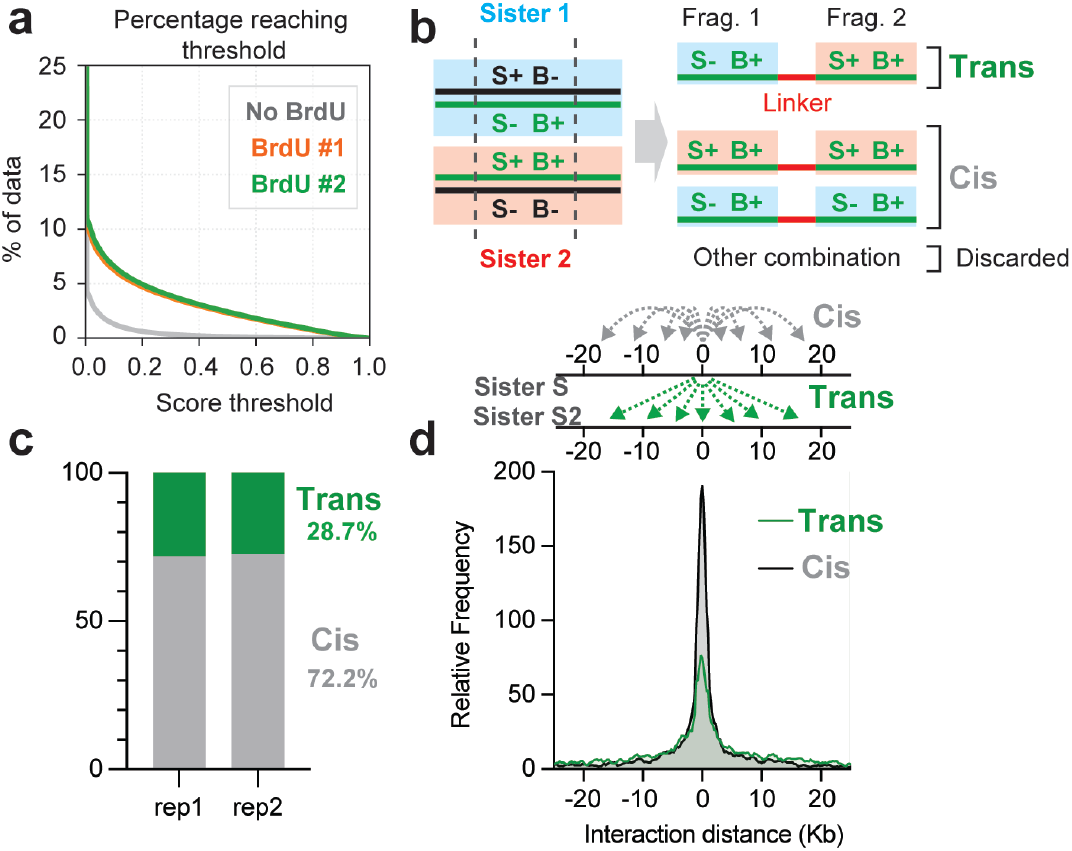
Analysis of trans-sister chromatin interaction by detection of BrdU. **a**) Percentage of reads that reach a BrdU score threshold between a control sample without BrdU and two samples incorporating BrdU. A threshold of 0.55 was selected for further analysis, this resulted in <5% false positive. **b**) Scheme of the selection process for the detection of trans-sister interactions, S denotes strand and B denotes BrdU. **c**) Graph showing the percentage of *cis* vs *trans* pairwise interactions **d**) The distribution of the genomic distance of trans-sister interactions (green) and cis-sister interactions (grey).

## DISCUSSION

The use of CAD notably improves ligation efficiency on crosslinked chromatin compared to MNase. We suspect that the longer extremities of linker DNA remaining after digestion is beneficial for subsequent end repair and ligation^31–33^. CAD digestion is easily optimized, and the sequenced data generates very homogeneous signal across the genome (Fig.S1a), suggesting minimal bias compared to MNase digestion. The high efficiency of DNA ligation allows several nucleosomes to be ligated together in long concatemers which are highly amenable for sequencing with long-read technology. A major benefit of this approach is that PCR amplification is not needed and so the resulting data is less subject to known biases introduced by amplification, moreover, direct sequencing also permits identification of base modification and nucleotide analogues.

CAD-C data does not provide clear evidence for predominant patterns of nucleosome folding such as the 30-nm fiber. If chromatin was organized as a stable zig-zag in some regions of the genome, ligation events from a nucleosome n could be expected to contact n+1 and n+2 with comparable probability, instead, CAD-C consistently shows a strong enrichment for n±1 over n±2 across different genomic contexts, arguing against a dominant, regular zig-zag connectivity – favoring a more linear organization in budding yeast. When focusing on the fraction of the long reads that ligate three consecutive mono-nucleosomes, patterns emerge that show similar frequency for linear (1-2-3) and intercalating (1-3-2) configurations. These configurations are compatible with sparse nucleosome stacking consistent with nucleosome clutches^28,43^. It is also important to consider whether proximity ligation approaches are well suited to detect differences in the interconnectivity between adjacent nucleosomes, indeed even in ideal structures with stacked or unstacked nucleosomes, the relative arrangement and proximity of linker DNA in the fiber is little changed, and it is perhaps unrealistic to expect that DNA ligase will preferentially ligate one connection over another. Thus, alternate approaches may ultimately be required to better understand how nucleosomes are arranged *in vivo*^43,44^.

Over long distances, the pairwise interaction maps produced by CAD-C are comparable in quality and resolution to the most advanced approaches currently available to date^22^. CAD-C matrices faithfully recapitulate all major chromatin organizational features that have been previously characterized through other methods. This includes the detection of Chromosomally Interacting Domains (CIDs) defined by insulator boundaries in G1 phase, as well as the formation of cohesin-mediated loops that become prominent in G2 phase. Importantly, CAD-C extends beyond simple pairwise interactions, as it is capable of detecting multiway chromatin contacts, exemplified by our identification of hubs formed by multiple cohesin-bound sites associating in G2. These results demonstrate that CAD-C represents a robust and reliable proximity ligation-based approach alternative to Micro-C with distinct benefits.

We present evidence that sister chromatids are near perfectly aligned in nocodazole arrested yeast cells a stage comparable with G2 phase in metazoa^45–47^. Given that CAD-C is performed in haploid yeast, such ligation products must occur by direct interactions of homologous sequences between two sister chromatids. Importantly, these interactions are abolished in G1 arrested cells or when cohesin is depleted in G2. Mapping the frequency of direct sister ligation across the genome produced a sister chromatid alignment score. This metric is inherently stringent: if sister chromatids are tightly aligned but offset by even one nucleosome, the score becomes zero. Consequently, this signal only measures simultaneous alignment and proximity and likely underestimates the true degree of sister chromatid connections. Remarkably, a centromeric nucleosome is as likely to ligate to its sister centromere in *trans* as it is to ligate to a neighbor nucleosome less than ∼20nm away in *cis*. This finding can only be explained if centromeric nucleosomes are closely paired in a majority of cells and may indicate a specific dimeric structure. Such physical interactions between sister centromeres appears to be mediated by cohesin (Fig.5b-d). Close coupling of kinetochores is suspected to occur in Meiosis-I where the monopolin complex bridges the two centromeres on sister chromatids and ensures they are segregated together^48^. The biological function of closely paired centromeres in G2 is not known, but may, in part, be promoted by the ability of inner kinetochore component Ndc10 to dimerize and interact with two distinct DNA molecules^49^, or through dimerization of partially wrapped centromeric hemisomes^50^.

Outside of centromeres, we find that sites of cohesin accumulation are also very closely aligned. At these places, direct ligation between homologous sequences in *trans* is less than that of centromeric sequences. Yet, given that ligation efficiency decays rapidly with genomic distance (Fig.2b-d) our data indicates that sisters are maintained with very close pairing. The contact frequencies we observe in *trans* are comparable to *cis* interactions between nucleosomes separated by less than 750bp i.e. N ligating to N+4 (Fig.2b-d and Fig.5s). Such contact frequencies could be explained if the two sisters are near perfectly aligned but are constrained by entrapment within a 30-40nm cohesin ring. We conclude that sister chromatids are frequently perfectly aligned at cohesin enrichment sites which are found every 5-15kb across the *S. cerevisiae*, genome^41^. Such pairing therefore ensures that the intervening DNA is also highly aligned between the two sisters, and this is evident in our data. Beyond cohesin peaks we find little evidence that sisters are preferentially cohesed at specific genomic locations, and we find no evidence increased cohesion at sites of replication termination (Fig.5c)^51^.

We utilized the ability of nanopore sequencing to detect BrdU incorporation and so resolve inter-sister interactions^42^. This approach is complementary to detecting homologous sequences in the same read but does not require perfect overlap between sisters to detect *trans* interactions. While detecting BrdU in very short nanopore reads is challenging, enough reads were successfully mapped to allow us to confirm that sister chromatids are near perfectly aligned across the genome. Future improvements will be driven primarily by BrdU/analogue detection (e.g., DNAscent) or by increased nanopore sensitivity itself. Continued development in either domain is likely to deliver finer sister-chromatid discrimination and lower false-negative rates that will empower CAD-C analysis even further.

Overall, our data significantly advances the understanding of cohesion and sister chromatid pairing. The near perfect alignment of sister chromatids we observe has previously been missed because the comparatively low resolution of Hi-C-based methods ^10,11^. Importantly, the long-range trans-interactions proposed in these Hi-C models can be explained by the interplay between loop-expanding cohesins and cohesive cohesins, which bind more stably to chromatin. Recent studies have shown that cohesive cohesins restrict loop expansion by non-acetylated cohesins^52^, resulting in the colocalization of cis-loop bases with cohesive cohesin complexes. The general inability of Hi-C to detect contacts <1.5 kb would mask short-range interactions while highlighting trans-interactions at the scale of loop sizes. Chromatin compaction through looping is revealed by the multi-way connectivity of ligated nucleosome chains captured in our single-molecule data. This connectivity pattern demonstrates that interaction hubs form at precisely paired sister chromatid loci, which simultaneously engage in long-range contacts with more distal genomic sites (Fig.5b, Fig.7).

**Figure 7:**
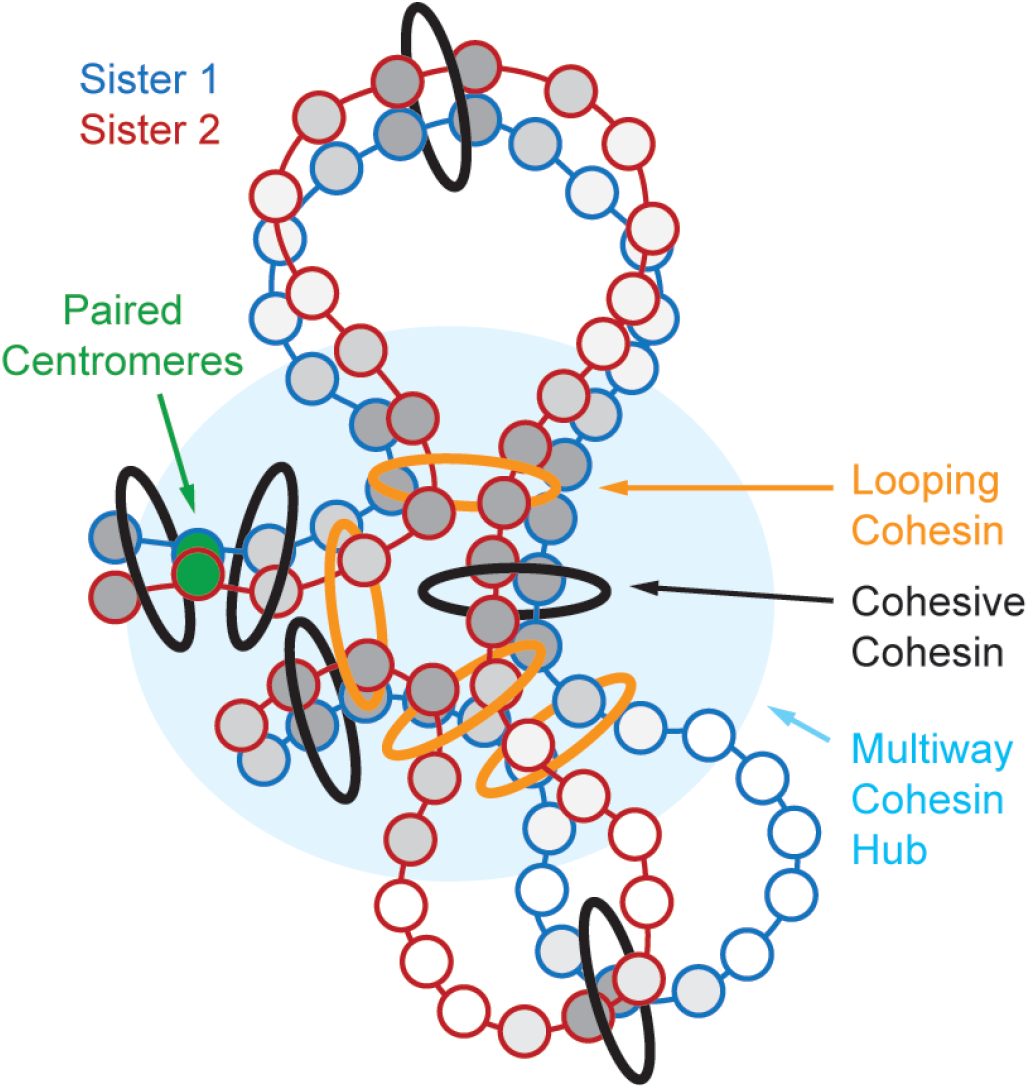
Model of sister chromatid cohesion: Proposed model in which sister chromatids are tightly aligned, with precise centromere alignment and pairing, frequent near perfect alignment at cohesin sites, and slightly looser alignment between cohesin sites. Increase in alignment degree is represented as darker shades of grey.

Cohesion establishment is tightly coupled to DNA replication through a sophisticated molecular mechanism. The acetylation of Smc3 at lysines 112 and 113 (K112, K113) by the acetyltransferase Eco1 stabilizes cohesin on DNA and renders it resistant to unloading pathway^53–55^ until cleavage by Separase in Anaphase^56,57^. Cohesion is established specifically during S phase^58,59^, at the replication fork^60^, where Eco1 is recruited through direct interactions with the replication processivity factor PCNA^61^. Importantly, the replisome can traverse through the cohesin ring^51,62^, allowing newly synthesized sister chromatids to be captured within the same cohesin complex as they emerge from the replication machinery. This intimate spatiotemporal coupling between replication fork progression and cohesion establishment provides an elegant mechanism to ensure that cohesin entraps precisely aligned, newly replicated sister chromatids. What is perhaps more surprising is that precise alignment persists long after replication has been completed. Sister alignment is clearly maintained at cohesin enrichment sites, whose locations in budding yeast are determined by patterns of convergent transcription^60^. Here transcription by RNA polymerase II sculpts cohesin localization and pushes cohesin along chromatin to preferred sites immediately prior to replication. Cohesion would then be established when the replisome passes through these sites. The near perfect alignment and close contact of sister chromatids may have distinct benefits for maintenance of genome integrity through DNA repair in G2 by homologous recombination^13,63^ and inheritance of gene expression patterns where epigenetic states may be copied between the perfectly aligned sisters. Self-reinforcing read-write mechanisms found in heterochromatin would presumably work across paired sisters^64,65^. Further, the tight and precise pairing of centromeres following DNA replication may facilitate the reestablishment of centromeres on the two daughter genomes, perhaps contributing to epigenetic inheritance^66,67^.

CAD-C should provide a valuable tool for studying chromosome structure and will be particularly useful for the investigation of how homologous sequences interact through cell division and in the maintenance of genome integrity.

## RESSOURCE AVAILABILITY

### Lead contacts

Axel Delamarre, Iestyn Whitehouse

### Data and code availability

Deposited data: Raw and process data are available at GEO: GSE312896

Code availability: Steps to follow to process raw data are available on Gihub:

https://github.com/Axel-Delamarre/Whitehouse-Lab/tree/main/Master_CAD-C

## ACKNOWLEDGMENTS

We thank Neeman Mohibullah and the IGO core at MSK for their help to sequence with nanopore. We thank Xiaolan Zhao and Armelle Lengronne for sharing yeast strains. We thank the Whitehouse and Remus lab for sharing reagents and materials. Work in the Whitehouse lab is supported by NIH grants GM152916 and GM129058. Research by MAB is supported by the Medical Sciences Research Council (MR/W031442/1). We also thank the Risca Lab, Rockefeller University, for sharing unpublished results regarding use of CAD for proximity ligation assays.

## EXPERIMENTAL MODEL AND STUDY PARTICICPANT DETAILS

### Yeast strains

yIW385: CEN.PK – MATa

x7319-c: MATa LEU2::BrdU-Inc URA3::GPD-TK7 pp5597: MATa HIS3::ADH1promoter-OsTIR1; SCC1-PK3-AID::Kan

## METHODS DETAILS

### Cell Cultures

75mL of 1×10^7^ cells/ml were grown in YPD and fixed in 1% formaldehyde for 15min at room temperature (RT). Formaldehyde was quenched with 0.75M Tris HCl, pH 8 for 1 minute. Cells were washed three times in 1X PBS at 4ºC. Cells were resuspended and flash frozen in 300μL of XLB-3 (Sorbitol 1M, NaCl 50mM, HEPES pH 7.5 10mM, MgCl_2_ 3mM). For culture in G1, cells were synchronized with α-factor at 2μg/mL for 3hr with re-addition of α-factor every hour before fixation. For culture in G2, cells were synchronized in G1 as above, released by filtration into YPD containing Nocodazole 15μg/mL (sigma, M1404) for 2h before fixation. For Auxin Induced Degradation, IAA (sigma, I3750) was added at 0.5mM. For BrdU incorporation, the yeast strain x7319-c was released from G1 synchronization in YPD supplemented with Nocodazole 15μg/mL and BrdU 400µg/mL (sigma B5002).

### CAD purification

pETDuet_huCAD_ICADL (Addgene, 100098) was transformed into T7 Express LysY/Iq competent cells (NEB, C3013I). Cells were grown in 2.4L of LB2YT (Tryptone 10 g/L, Yeast Extract 10 g/L, NaCl 5 g/L) until OD reached 0.38 and induced with IPTG 0.5mM for 4h at 37°C. Cells were pelleted at 4000 x g for 30min, and the pellet was stored at −80C. The pellet was resuspended in 70 mL Lysis buffer (NaCl 300 mM, HEPES pH 7.5 50 mM, Glycerol 10% w/v, Imidazole 20 mM, 1X Protease Inhibitor (Pierce, A32963) and sonicated on ice for 10 seconds on, 20 seconds off at 50% amplitude for 6 min. The sonicated mix is transferred to 2 × 40 mL high speed centrifuge tubes, and spun down at 18,000 x g for 1 hour at 4° C. The supernatant was recovered and passed through a HisTrap FF (Cytvia, 17528601). The column was washed with 25 mL of Wash Buffer (NaCl 300 mM, HEPES pH 7.5 50 mM, Glycerol 10% w/v, Imidazole 40 mM, 1 x Pierce Protease Inhibitor Cocktail Tablet). The complex was eluted with 6mL of Elution Buffer (NaCl 0.3M, HEPES pH 7.5 50 mM, Glycerol, 10% w/v, Imidazole 0.5 M) and stored at −80°C. The purified CAD/ICAD was dialyzed into Exchange Buffer (NaCl 400 mM, HEPES pH 7.5 20 mM, Glycerol 10% w/v, DTT 5mM) and aliquoted in 100μL containing 50μg of protein. On the day of the experiment, 87 μL of CAD was activated by addition of 10μL of TEV buffer and 3μL of TEV enzyme for 1hr at 30°C (NEB, P8112S).

### Linker annealing

Top linker: /5phos/GTCGCCTACCTAC and Bottom linker: CTTTTT/ideoxyU/GCGCGTAGGTAGGCGACT, were mixed at 25μM in Annealing Buffer (100 mM KAc; 30 mM HEPES pH 7.5). Annealing was performed by heating to 94ºC for 4 minutes and letting the heat block cool down to RT overnight.

### CAD-C method

Cells were broken using freezer mill (Spex SamplePrep,6875D) for 5 cycles, 2 min, 15cps. After verification of breakage under the microscope, cells were resuspended in 900μL of XLB3 (Sorbitol 1M, NaCl 50mM, HEPES pH 7.5 10mM, MgCl_2_ 3mM, supplemented with protease inhibitors, A32955, Pierce) and cross-linked by addition of DSG at 3mM final (Disuccinimidyl glutarate, A35392, ThermoFisher) for 30 minutes at 30°C. DSG was quenched by addition of 100μL of 1M Tris HCl, pH 8 for 1 minute. Cells were washed 3 times in HS buffer (Sorbitol 1M, HEPES pH 7.5 10mM), resuspended in CAD buffer (Sorbitol 1M, HEPES pH 7.5 10mM, DTT 5mM, IGEPAL 0.1%, MgCl_2_ 1mM, protease inhibitors A32955, Pierce) and supplemented with pre-activated CAD. Chromatin was digested for 1hr at 37ºC with 1000 RPM agitation. Cells were placed on ice for 2 min and washed twice in SB buffer (10mM HEPES pH 7.5, 0.1% IGEPAL, 1mM EDTA, 1mM DTT) with pelleting between washes at 3000xG before resuspension in 500μL of SB buffer.

End-Repair and A-tailing were performed with NEBNext Ultra II DNA library Prep Kit (NEB, E7645) for 30min at 20ºC with constant mixing, then 30min 65ºC with constant mixing. The chromatin was incubated for 2 min on ice, washed twice in 500 μl of SB buffer. The chromatin was resuspended 225 μL of HEPES pH 7.5 10mM, supplemented with 25µL of annealed linker at 25μM (top linker: /5phos/GTCGCCTACCTAC, bottom linker: CTTTTT/ideoxyU/GCGCGTAGGTAGGCGACT. Linkers ordered from IDT) and 250μL Blunt/TA ligase (NEB, M0367L) for 40 minutes at RT. The chromatin was washed three times in 500μl of SB and resuspended in 50µL of rCutsmart buffer 1X (NEB, B6004). 5µL of USER enzyme (NEB, M5505) was added and the reaction was incubated for 15 min at 37ºC. The reaction was cooled to RT and 200ul of HEPES buffer 10mM was added followed by 250uL of Blunt/TA ligase MM 2x. The ligation was incubated for 1hr at RT. The reaction was washed once in 500μl of SB buffer and resuspended in 100µL of decrosslinking buffer (100mM Tris HCl pH 7.5, 5μL of RNAse cocktail (Invitrogen, AM2286), 5µL of Proteinase K 20mg/mL) overnight at 55°C. The DNA was purified using a Zymo column (Zymo Research, D4014). The nanopore library was produced using the kit SQK-LSK114. Each library was sequenced on one Promethion.

## QUANTIFICATION AND STATISTICAL ANALYSIS

### Data availability

Raw and processed data are available on GEO with accession number: GSE312896

### Basic data processing

Data processing is described in detail on GitHub: https://github.com/Axel-Delamarre/Whitehouse-Lab/tree/main/Master_CAD-C. Briefly, the fastq are segmented based on the identification of the linker sequence and aligned on the genome using minimap2. Reads with mapq score below 40 were filtered out as well as reads with a hairpin structure (symmetrical and containing fragments with the same arrangement and coordinates but opposite strands). We believe that hairpin reads are generated during the end-repair and ligation steps where the 3’end of one strand ligates to the 5’ end of the opposite strand. Each fragment is then reported in a *read_info*.*txt* file that contains the following columns: chromosome, start, end, fragment size, midpoint, strand and read-ID.

### Generation of high-resolution pairwise interaction matrices

Reads are then grouped based on their read-ID, creating multiway tag groups. All the unique pairwise combinations of a given group are reported in a table that is then used to produce interaction matrices (midpoint-to-midpoint) using juicer-tools to produce *.hic* maps and cooler to produce *.mcool* map. For this analysis only, fragments longer that 260bp, the midpoint of the first nucleosome is calculated by adding 83 to the starting point coordinate.

### Matrix Balancing

Matrix balancing is a common practice in analyzing 3C data to normalize data, ensuring that each row and column sum to the same value. This approach helps to mitigate genome coverage and digestion biases, providing a more uniform representation of each restriction fragment. However, in technologies like Micro-C and CAD-C, which essentially map individual nucleosomes, natural biological variations in nucleosome occupancy such as nucleosome-free regions (NFRs) that typically have lower signals become significant. Balancing in these contexts can introduce artificial signal which confounds interpretation. Further, given that the read coverage across the genome generated by PCP shows minimal bias, the interaction matrices in this article are not balanced.

### Precise interactions distances. (Fig.2)

Only mono-nucleosome reads are considered, with fragments from 110 to 190 bp. Interestingly, a notable fraction of reads shows an interaction distance of –1bp when considering extremities to extremities. Inspection of long reads shows these fragments are separated by a linker and are bona fide proximity-ligation products. This can happen if CAD leaves a 1-nt 5′ overhang that is filled during end-polishing on both extremities before linker ligation and subsequent proximity ligation.

### Sister interactions by overlapping fragments from the same long-read. (Fig.5)

Pairwise tables generated directly from the *read_info*.*txt* are used to determine overlap between fragments sharing the same read-ID and chromosomes. Prefect overlaps (i.e. fragments with exactly the same coordinates), or fragments that overlap by less than 15 bp, or fragments shorter than 100bp are filtered out and not included in the analysis. The total overlap is then computed for each base in the overlapping read and summed across the genome to generate a bigwig track. The multi-way interaction matrix in Fig 5b was generated by extracting only read-IDs with overlapping fragments (i.e. *trans* interactions), pairwise interactions were then calculated from these reads to generate a matrix.

For quantifications of trans (n to n trans) vs cis (n to n+1), we only considered fragments directly ligated to each other within a long read. For an accurate measure we only considered mono-nucleosome size fragments (140bp-200bp). We took as input the total number of interactions between mono-nucleosomes, we counted as *trans* the number of overlapping reads (overlap >30bp). We counted as *cis* (n to n+1) when midpoint interactions distances are between 150 and 200bp.

### BrdU detection (Fig.6)

All pod5 files were converted into Fastq files and split according to the linker sequence. All reads were aligned to the reference genome with Dorado (Oxford Nanopore Technologies) and the resulting bam file was sorted and indexed. BrdU containing reads were called with DNAscent detect v4.1.1 ^68^ using the relaxLength branch settings with a quality control threshold on the adaptive banded signal to basecall alignment of −6 and a minimal length of 100 bp. A BrdU score was calculated for each read which represents the fraction of BrdU positive positions (BrdU score ≥ 0.55) across the entire read. Only reads with a BrdU score that satisfy this threshold were used for subsequent analysis. Fragment BrdU score obtained this way are assigned to the reads processed in the main workflow. Pairwise tables are then processed and only BrdU positive reads are analyzed. Pairs of reads sharing the same chromosome are annotated *cis* interactions if they share the same strand and *trans* if they have opposite strands. Midpoint distance between the two members of the pair is reported.

Plots and statistical analysis were performed using GraphPad prism (v10) and are detailed in the figure legends. Error bars represent SD unless stated otherwise.

## SUPPLEMENTAL FIGURES

**Supplemental Figure 1:**
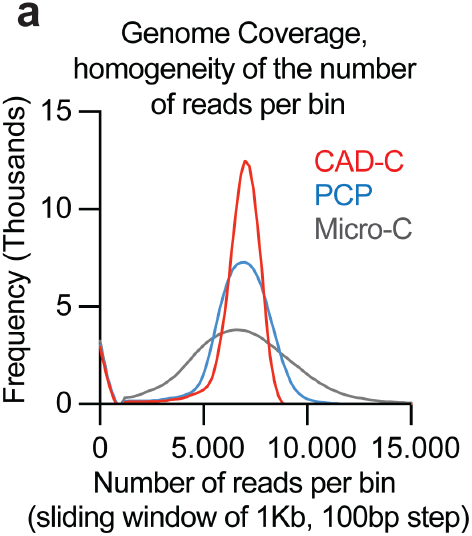
Read coverage of PCP compared to Micro-C (Costantino et al. 2020) and PCP (Delamarre et al. 2025) as a number of reads per sliding windows of 1kb with 100bp steps.

**Supplemental Figure 2:**
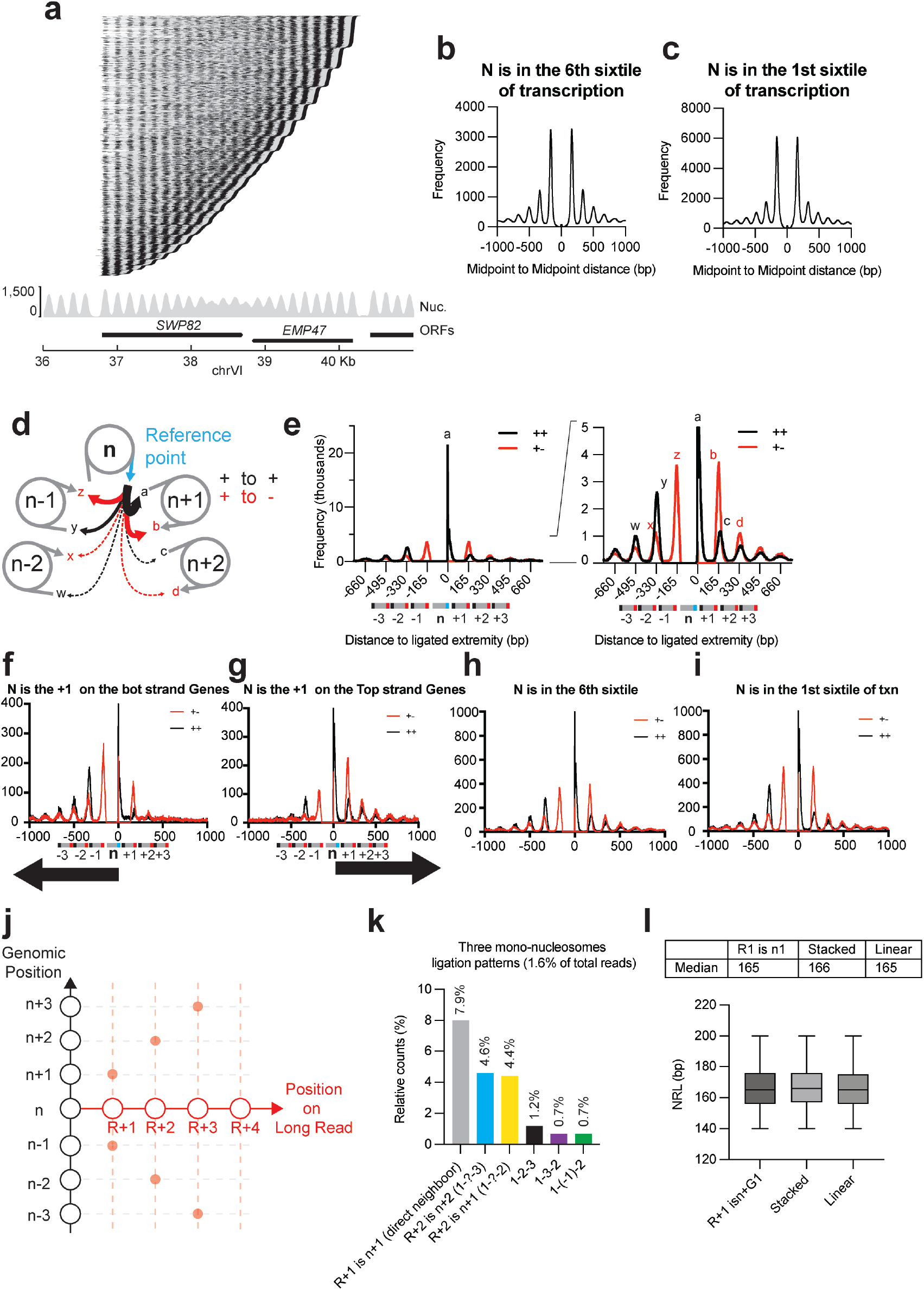
a) Long read fragment composition over the convergent genes *SWP82* and *EMP47* in G1, each line corresponds to a long read and each black dot to a ligated chromatin fragment. b) Midpoint to midpoint distance ligation distribution on the 6^th^ sextile of transcribed genes (least transcribed). c) Midpoint to midpoint distance ligation distribution on the 1^st^ sextile of transcribed genes (most transcribed). d) Scheme of the extremities-to-extremities ligation distribution. e) Extremities to extremities ligation distribution, letters showing specific ligations are shown in d; + to – in red and + to + in black. f) to i) Extremities to extremities ligation distribution. + to – in red and + to + in black. Around the +1 nucleosome of bottom strand genes f) or top strand gene g). On the 6^th^ sextile of transcribed genes (least transcribed) h) or 1^st^ sextile (most transcribed) i). j) Diagram explaining the heatmap in Fig2.e measuring the frequency of ligation between fragments on long reads. The relative position on the long read is plotted on the X-axis (R, R+1, etc…) and the relative position on the genome is plotted on the Y-axis (n, n+1, etc..). k) Quantification of the signal of the heatmap in Fig2.e i) Nucleosome Repeat Length (NRL) of the configuration identified in Fig2.g) showing no differences between neighbor nucleosome ligation, stacked configuration and linear configuration.

**Supplemental Figure 3:**
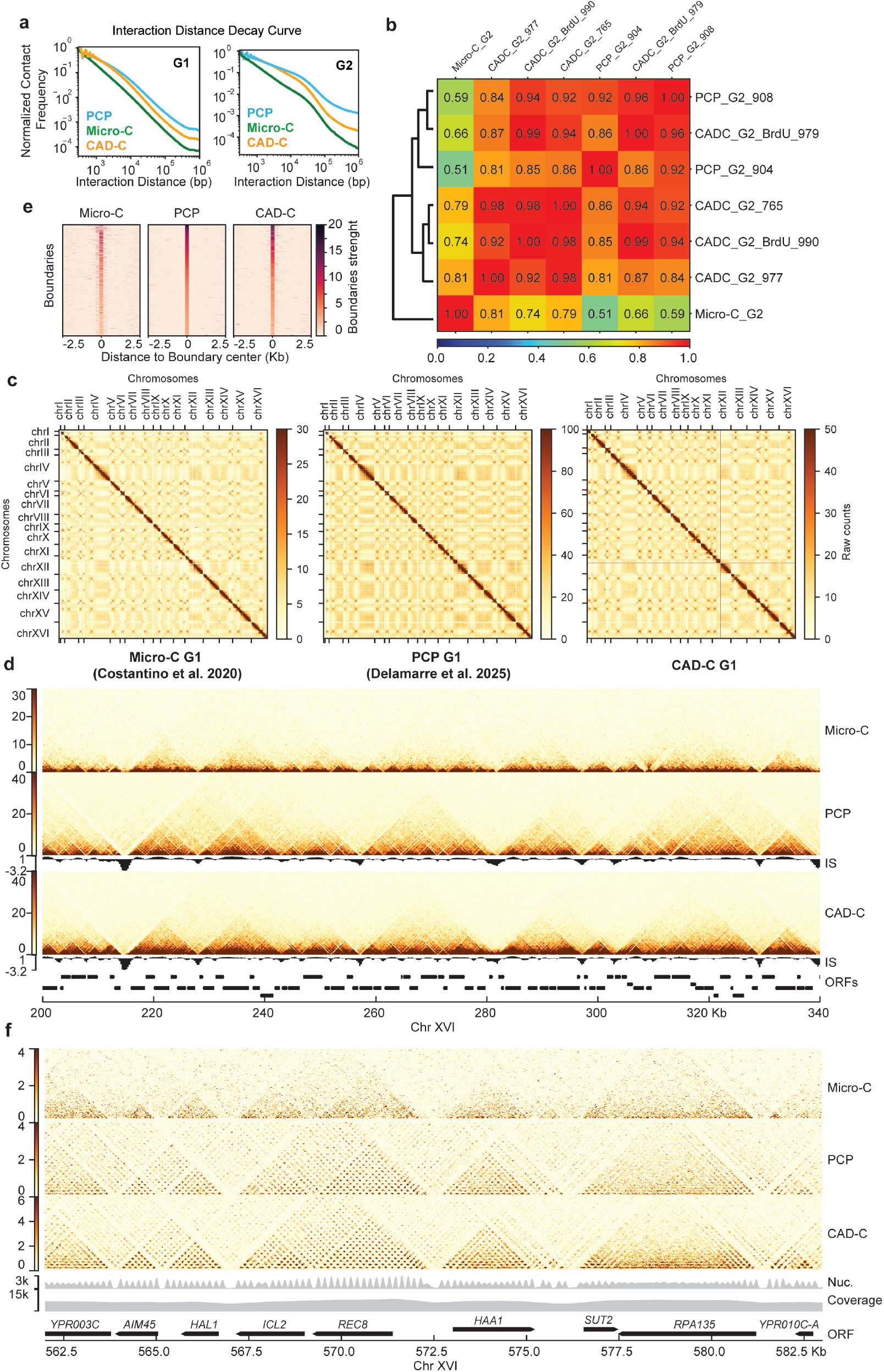
a) Interaction distance decay of Micro-C (Costantino et al. 2020), PCP (Delamarre et al. 2025) and CAD-C (this paper) in G1 and G2. b) Pearson Correlation of interaction matrices of Micro-C (Costantino et al. 2020), PCP (Delamarre et al. 2025) and CAD-C (this paper) in G1 and G2. c) Comparison of genome wide matrices produced by Micro-C (Costantino et al. 2020), PCP (Delamarre et al. 2025) and CAD-C (this paper) in G1. Note the high density of interactions between the centromeres of different chromosomes. d) 20Kb scale comparison of genome wide matrices produced by Micro-C, PCP and CAD-C in G1. e) Heatmap of insulation scores of boundaries of Micro-C (Costantino et al. 2020), PCP (Delamarre et al. 2025) and CAD-C (this paper) in G1. Sorted on the PCP data. f) 2Kb scale comparison of genome wide matrices produced by Micro-C, PCP and CAD-C in G1.

**Supplemental Figure 4:**
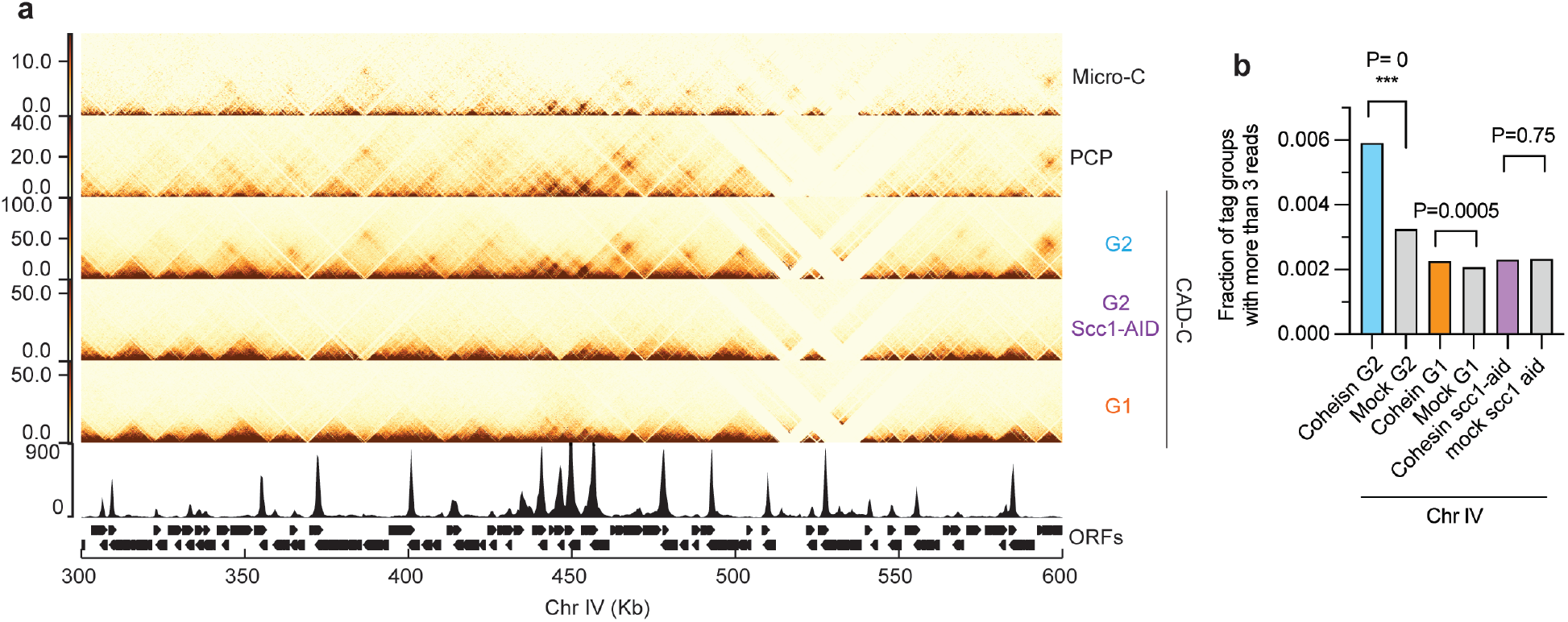
a) Comparison of genome wide matrices produced by Micro-C, PCP, WT CAD-C and under cohesin depletion (Scc1-AID) in G2 (Nocodazole arrest) b) Quantification of the frequency of multiway interaction with more than three fragments in different cohesin accumulation sites in G1, in G2 or in G2 after cohesin depletion.

**Supplemental Figure 5:**
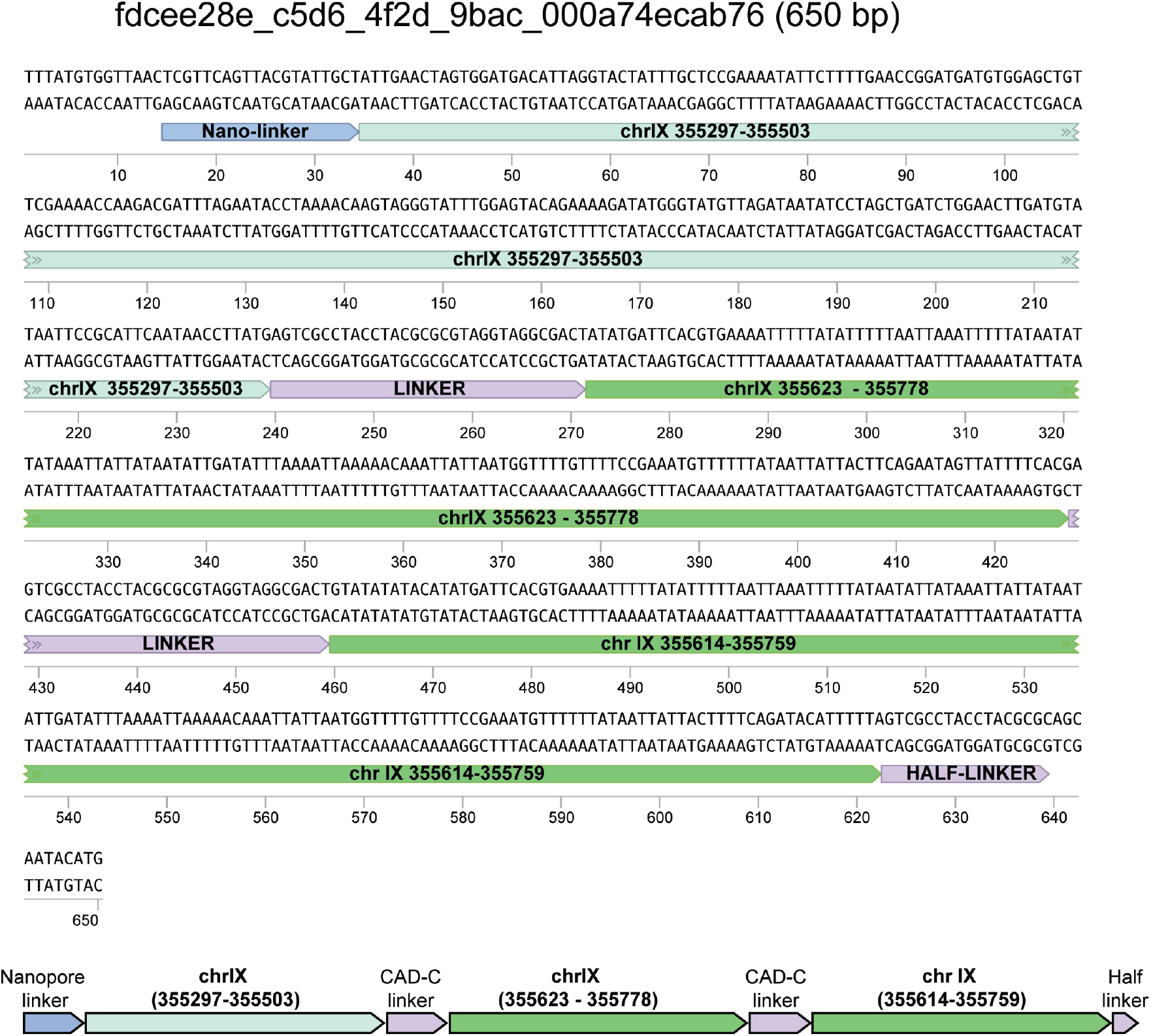
Example of a long-read containing the same segment twice (dark green) covering the centromere of ChrIX.

**Supplemental Figure 6:**
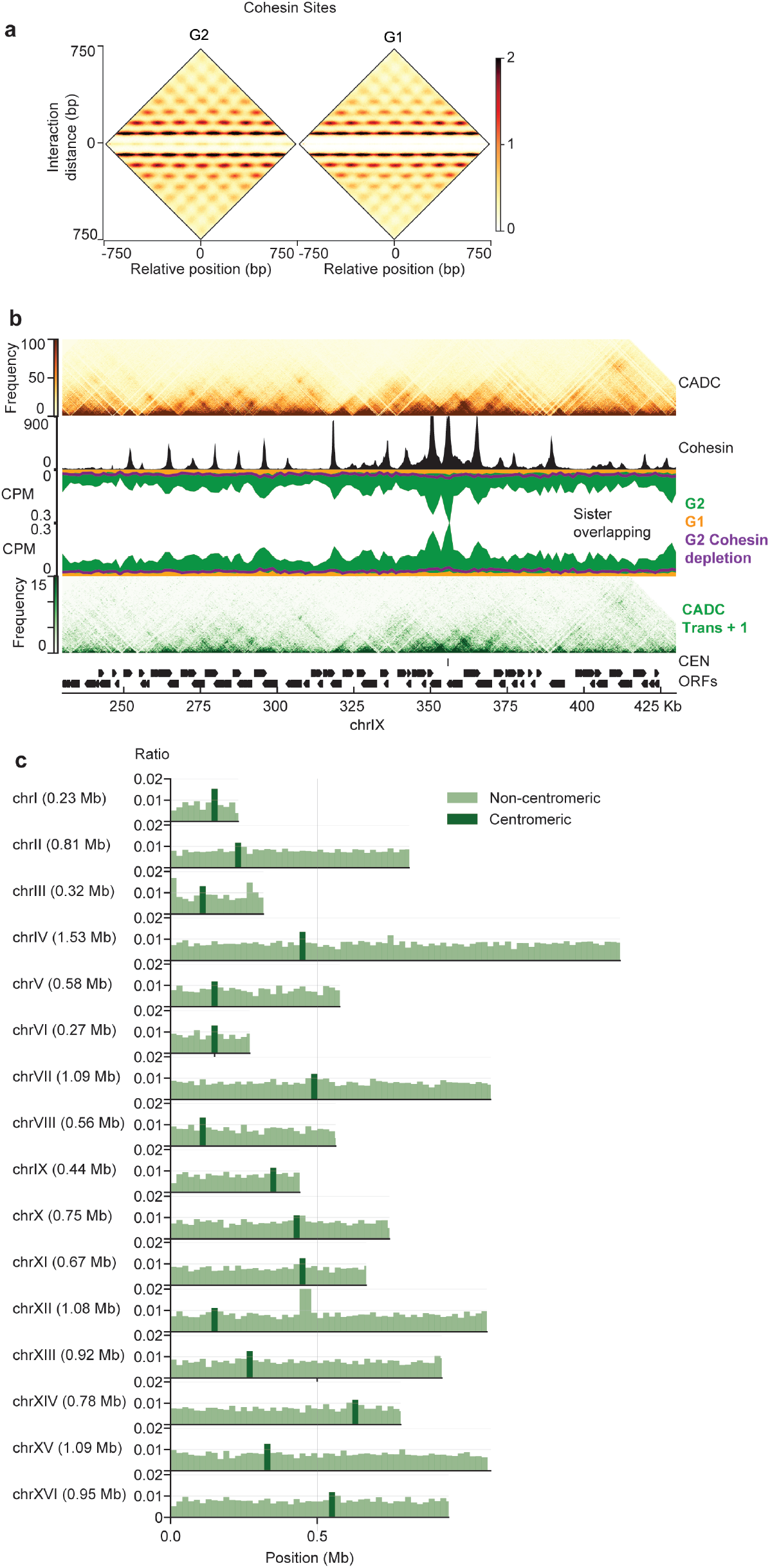
a) Pile-up analysis of CAD-C maps in G2 and G1 over cohesin accumulation site. A clustering of all the cohesin sites was first conducted without supervision to allow alignment according to nucleosome positions. b) Chromatin landscape representing interaction matrix (top panel); cohesin ChIP dataset (black) (Costantino *et al*. 2020); mirrored sister chromatin ligation scores in G2 (green), G1 (orange) and after cohesin depletion in G2 (purple); interaction matrices based on long reads that contain sister chromatid ligation fragments as well as long-range interactions (lower panel). c) Siter nucleosome ligation scores over the 20Kb windows, as a ratio of number of overlaps over total number of reads on the whole genome of *S. cerevisiae*. Bins containing the centromere are dark green.

**Supplemental Figure 7:**
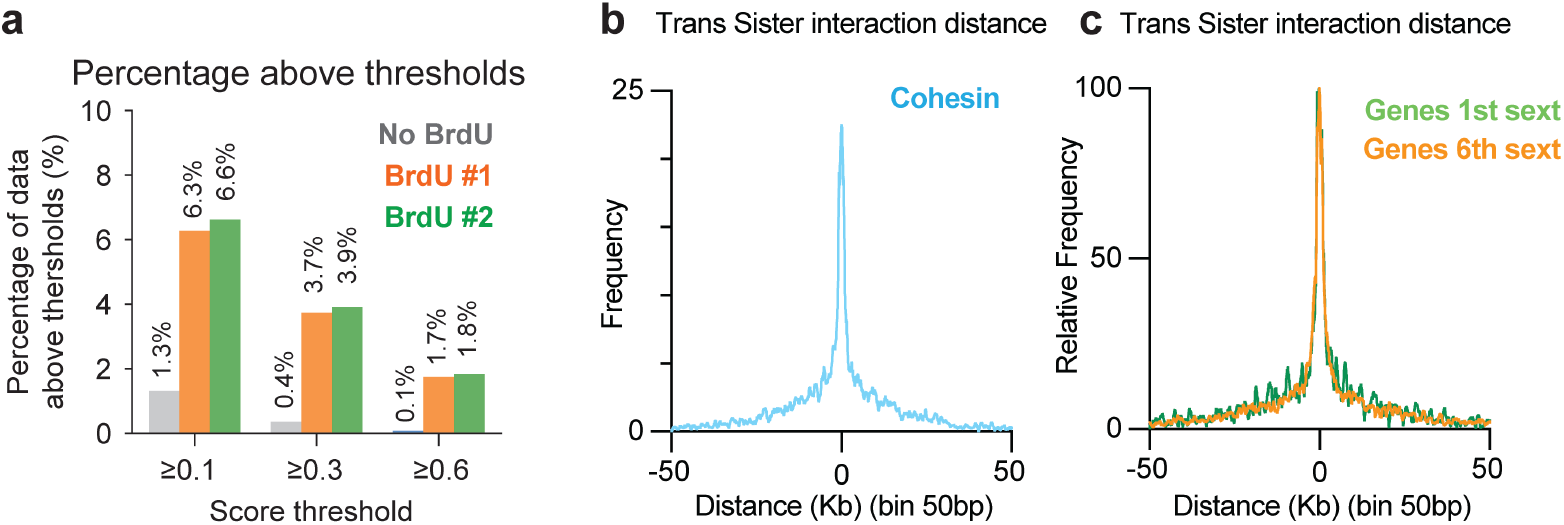
a) Percentage of reads that reach a BrdU score threshold between a control sample without BrdU and two samples incorporating BrdU. b) Distance between the two fragments of a *trans* interaction distance around cohesin peaks. (A threshold of 0.3 was use for this analysis) c) Distance between the two fragments of a *trans* interaction distance over the most transcribed genes (1 sextile, green) and the least transcribed genes (6^th^ sextile, orange). (A threshold of 0.3 was use for this analysis)

## Notes

### Competing Interest Statement

The authors have declared no competing interest.

### Summary of Updates

Quantification in figures 4 and 5 were updated.

